# Weber’s Law in walking: sensory scaling is observed in multi-sensory, dynamic tasks

**DOI:** 10.64898/2025.12.01.691589

**Authors:** Marcela Gonzalez-Rubio, Pablo Iturralde, Gelsy Torres-Oviedo

**Affiliations:** Department of Bioengineering, University of Pittsburgh, Pittsburgh, PA, United States; Departmento de Ingeniería, Universidad Católica del Uruguay, Montevideo, Uruguay; Center for the Neural Basis of Cognition, University of Pittsburgh and Carnegie Mellon University, Pittsburgh, PA, United States

## Abstract

We frequently adjust our behaviors in response to environmental changes. This behavioral flexibility requires adequate sensitivity to external stimuli to maintain optimal motor performance under evolving task demands. There is abundant empirical evidence that sensitivity scaling follows Weber’s Law. Weber’s Law states that the perception of a sensory stimulus is scaled by the magnitude of the background sensory context. However, Weber’s Law has so far been assessed only in uni-sensory static tasks, and it remains an open question whether this principle extends to multi-sensory, dynamic motor tasks. To address this question, we assessed somatosensory perception of relative leg motion (i.e., speed differences between right and left legs) in healthy young adults. We hypothesized that sensitivity to differences in leg speed would follow Weber’s Law. We estimated participants’ sensitivity to speed differences (sensory stimuli) using two-alternative forced choice (2AFC) tasks. Participants walked at a testing speed representing distinct sensory contexts: slow speed (low-intensity sensory context), comfortable speed (medium-intensity sensory context), and fast speed (high-intensity sensory context). All groups compared their assigned testing speed against a common reference speed. We found that sensitivity to speed differences followed Weber’s Law at both slow and fast non-habitual walking speeds, but this scaling differed when walking near the comfortable walking speed. Moreover, Weber’s Law scaling was reproduced by a drift–diffusion model that used only reaction times to quantify sensitivity, indicating that the evidence accumulation process described by the model can account for Weber’s Law scaling in multi-sensory, dynamic motor tasks, as well as in static, uni-sensory tasks.

**Author summary:** Adapting motor behaviors to environmental changes requires precise sensory perception to maintain optimal performance. Weber’s Law, a fundamental principle of psychophysics, describes how sensory perception is scaled relative to background stimulus intensity to prevent sensory overload; however, its applicability to multi-sensory dynamic motor behaviors like locomotion remains unclear. Our results show that Weber’s Law partially explains how we perceive leg movement differences during walking. We found that sensitivity to speed differences scales following Weber’s Law when walking at non-habitual slow or fast speeds. However, near comfortable walking, people are actually more sensitive than at other speeds, departing from Weber’s Law. A drift-diffusion model replicated Weber’s Law scaling during walking, showing that the same mechanistic framework can explain the scaling of sensation in both static, uni-sensory tasks and dynamic, multi-sensory activities like walking.

## Introduction

Successful motor control in dynamic environments depends on the continuous detection, integration, and scaling of sensory information across multiple modalities. Tasks such as reaching or walking require the nervous system to combine proprioceptive, tactile, visual, and vestibular cues to perceive and respond to changes in movement dynamics. The ability to detect differences in movement parameters, such as changes in limb speed, force, or movement trajectory, is critical for maintaining stable and coordinated behaviors. Previous work has characterized the motor responses involved in adaptation across various motor tasks [1–4], yet the perceptual principles governing how humans detect changes in expected movements remain poorly understood. In particular, little is known about how sensory scaling operates in multi-sensory, dynamic motor contexts.

Weber’s Law provides a framework for understanding how perceptual sensitivity scales with stimulus intensity. It states that the smallest detectable difference between two stimuli is proportional to the magnitude of the environment-specific sensory signals. In other words, humans detect differences as proportions rather than absolute values [5]. For example, adding 10 pounds to a 10-pound weight is easily noticeable, whereas the same 10-pound difference is barely perceptible when added to a 100-pound weight. This principle has been consistently demonstrated in static, uni-sensory tasks across sensory modalities. In vision, observers maintain constant discrimination for random dot motion across different signal-to-noise ratios [6]. The same scaling has been observed across a variety of perceptual tasks, such as pure tone intensity discrimination [7, 8], vibrotactile discrimination [9], joint position (proprioception) [10], and smell [11]. This cross-modal consistency suggests Weber’s Law reflects a fundamental organizational principle of sensory systems.

Whether Weber’s Law applies to multi-sensory dynamical motor tasks remains an open question, and walking provides an avenue to examine this. Walking is a whole body movement that requires continuous integration of proprioceptive and cutaneous sensory feedback, vestibular information, and efferent copies of motor commands to detect changes in the environment that require gait adjustments [12–20]. Detecting subtle differences in walking speed between legs is essential for maintaining stability and coordinated movements while adapting to different terrains. Walking then can be used as a model for evaluating whether the principles of sensory scaling described by Weber’s Law extend to the multi-sensory domain of dynamic and active sensory processing during whole-body movement.

Reaction times, defined as the time interval between stimulus onset and people’s response to it, provide a complementary measure of the temporal dynamics underlying perceptual judgments. Drift diffusion models (DDMs) describe this decision-making process by modeling perceptual discrimination as the accumulation of noisy sensory evidence over time until a decision is reached, with the accumulation rate influenced by stimulus magnitude [21–23]. Prior research in the auditory domain has demonstrated that timing patterns are related to sensory stimulus scaling, such that higher baseline stimulus magnitudes systematically slow the rate of evidence accumulation, resulting in decreased sensory sensitivity to the same stimulus difference [8]. From these findings, we asked whether DDM-based analyses of reaction times during walking speed discrimination could also be used to reveal changes in perceptual sensitivity, providing insight into whether the perceptual computations governing multi-sensory motor behaviors, such as walking, share common principles with those observed in uni-sensory static tasks.

Our primary objective was to determine whether perceptual sensitivity to speed differences during walking follows Weber scaling. If Weber’s Law applies to walking speed perception, we expect that individuals will be more sensitive to speed differences when walking slowly than when walking quickly. In other words, sensitivity should decrease linearly as the mean walking speed increases, so that the sensitivity relative to the mean walking speed, known as the Weber fraction, remains constant across all walking speeds. Additionally, we tested if the evidence accumulation described by drift diffusion models fitted solely to reaction times could reproduce said scaling. Understanding these perceptual mechanisms has broader implications for theories of sensorimotor control and could inform the development of targeted rehabilitation interventions by identifying walking speed conditions where individuals show optimal sensitivity to gait asymmetries.

## Results

We characterized perceptual sensitivity to speed differences in thirty-nine neurotypical young adults walking at distinct background speeds: 0.7 m/s (slow), 1.05 m/s (reference), 1.4 m/s (comfortable), and 1.75 m/s (fast). To avoid fatigue, each participant completed two conditions (Fig.1): a test speed (slow, comfortable, or fast), randomly assigned, and the reference speed, which is within the range of habitual speeds in young people [24]. At each speed, participants performed two-alternative forced-choice (2AFC) perceptual tasks in which a speed difference was introduced between the legs (Δ*V* = *V*_*R*_ − *V*_*L*_) while holding the mean speed across belts at the specified background speed (Fig. 1B). Participants indicated which leg felt slower using handheld controllers.

**Fig 1.**
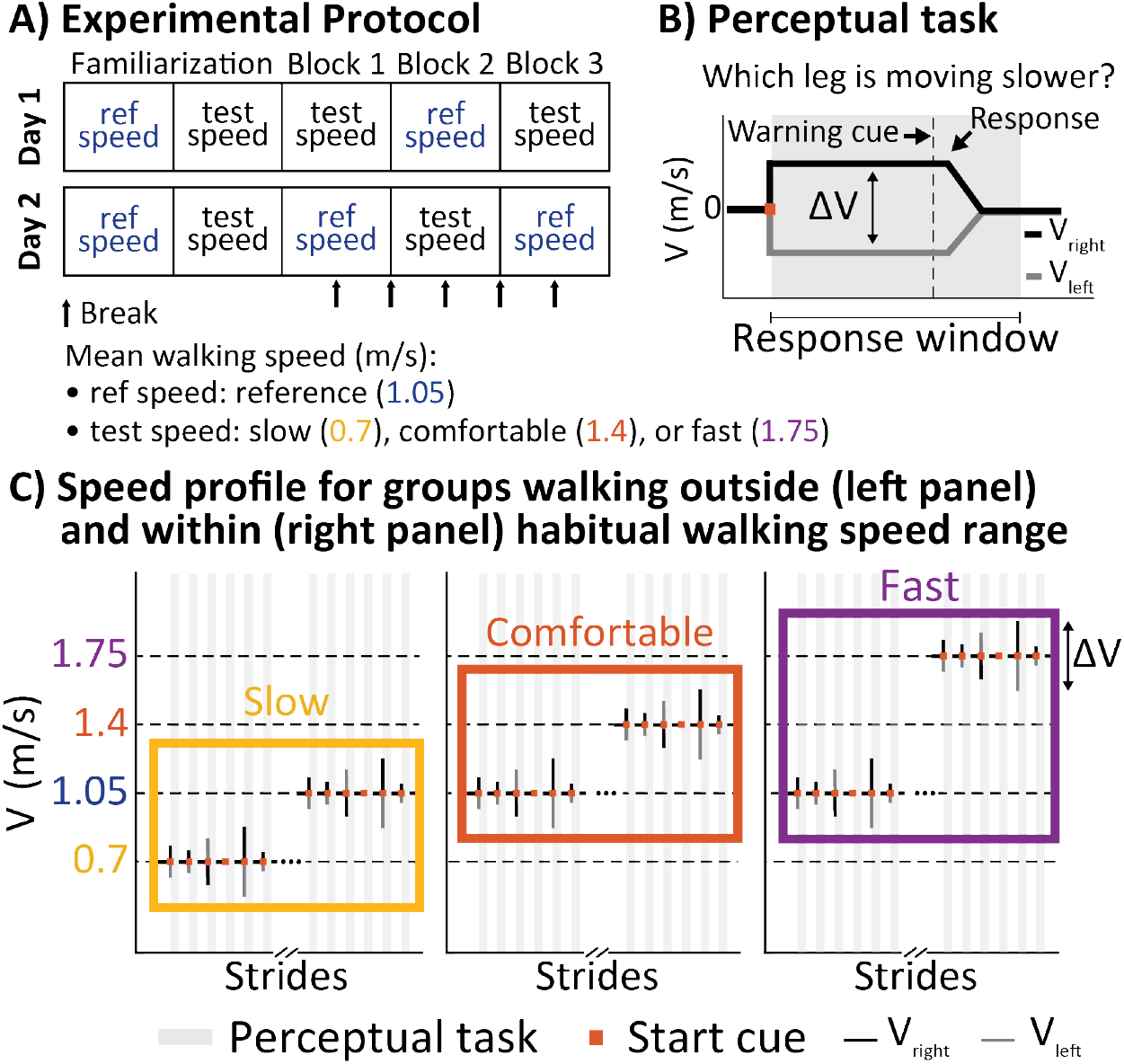
**A)** Schematic of the experimental protocol. Participants completed two data collection sessions on separate days (i.e., one day apart on average). Each participant experienced two distinct average walking speeds between their legs: a “reference” speed (ref in blue) and a “testing” speed (test in black). The reference speed was always 1.05 m/s, while testing speeds were either slow (0.7 m/s), comfortable (1.4 m/s), or fast (1.75 m/s) speeds for healthy young adults. Participants performed 3 data collection blocks. The speed conditions were interleaved within each block, and the first speed experienced was randomized across participants to counterbalance any potential effects of encountering a slower or faster speed first. The treadmill was stopped between blocks and once within each block to allow for 3-minute breaks, minimizing mental and physical fatigue (arrows indicate where the breaks occurred in the schematic). **B)** Perceptual 2AFC task schematic. The task began with participants walking with both legs moving at the same speed, followed by an abrupt transition to a belt speed difference and a start cue (red square) indicating the beginning of the response window (gray shaded area). The response window lasted 8 strides for the Slow and Comfortable group and 9 seconds for the Faster group. The perceptual task ended once a response was recorded or if the participant failed to respond by the end of the window. **C)** Example belt speed profiles during each speed condition. The left and right belts are shown as vertical gray and black bars, respectively. Each box represents a group: Slow (yellow, 0.7 m/s), Comfortable (orange, 1.4 m/s), and Fast (purple, 1.75 m/s). All groups shared the reference speed of 1.05 m/s. Each group of participants experienced walking at their group-specific testing condition and the reference condition while performing the perceptual tasks. Stimulus magnitude presentations were pseudo-randomized.

### Perceptual sensitivity to speed differences scales with walking speed

Psychometric functions describing participants’ responses in the 2AFC tasks are shown in Fig. 2A. These logistic fits, defined as *p*(choice = “left is slower”) 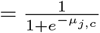, with 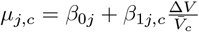, were computed for each participant (*j*) and speed condition (*c*), using a Generalized Linear Mixed-Effects Model applied to the complete dataset (see Methods). In line with Weber’s Law, the psychometric curves for participants walking at the testing and reference conditions overlapped when plotted against relative speed differences (top panels). Consistently, the same data plotted against absolute speed differences (bottom panels) show that slopes decreased with walking speed, being steeper at slower speeds (yellow) and shallower at faster speeds (purple). Taken together, these qualitative patterns support Weber’s prediction that relative rather than absolute sensitivity to speed differences is preserved at distinct background speeds.

**Fig 2.**
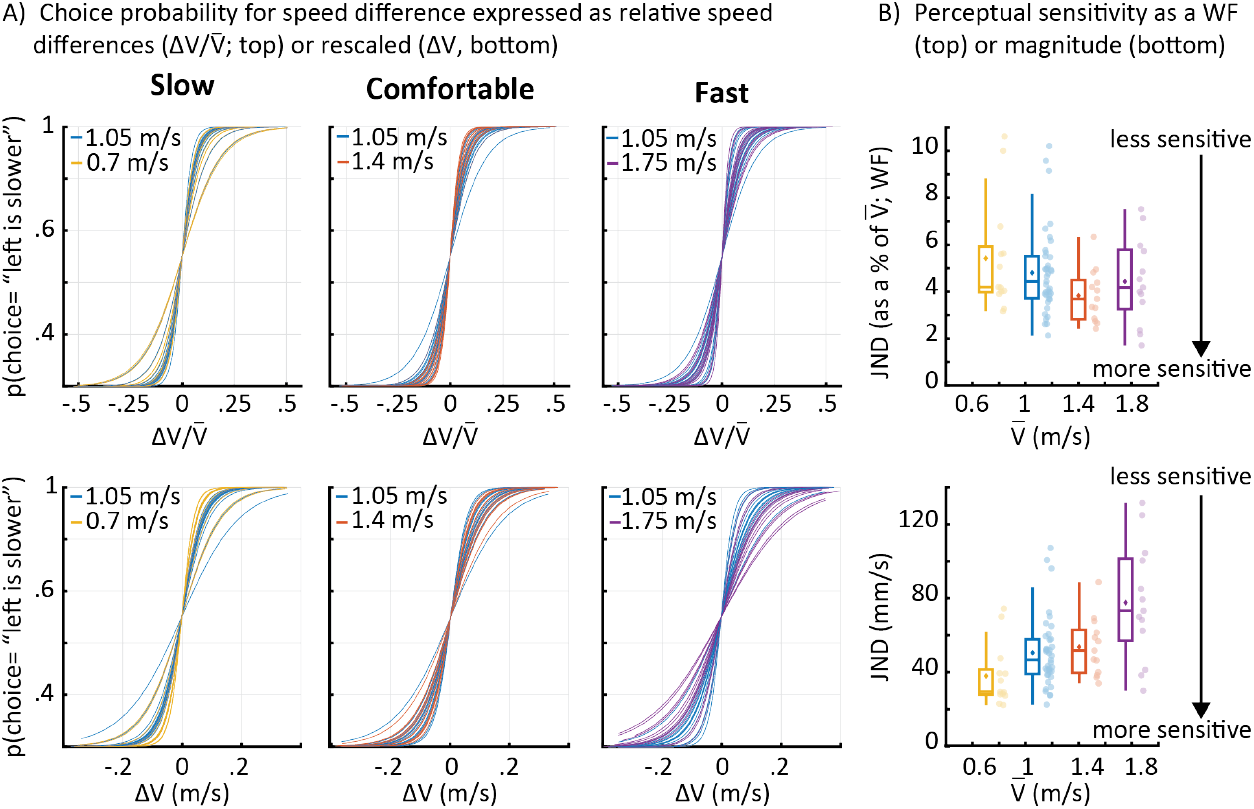
Sensitivity analysis across walking speeds. **A)** Psychometric functions resulting from fitting choice data as a function of the relative speed difference 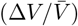 for the three groups (top panels, left to right), each tested under two speed conditions. Thin lines represent individual participant fits. Colors denote the testing condition: slow (0.7 m/s) in yellow, reference (1.05 m/s) in blue, comfortable (1.4 m/s) in orange, and fast (1.75 m/s) in purple. Under Weber’ s Law, psychometric functions for different speeds would overlap. Bottom panels show the same data as a function of the absolute speed differences (Δ*V*). Under this testing we would expect differences in the slopes of the psychometric functions across mean walking speeds. **B)** Just Noticeable Differences (JNDs) across the walking speed continuum show Weber’s Law. Reference speed information was pooled across all groups. The x-axis represents the average walking speed, and the y-axis shows the JND, where lower values reflect greater sensitivity to speed differences. For each tested speed, the distribution of JNDs is shown using box plots (median, interquartile range), means are represented inside the box plots as a diamond, and individual participant data are represented as colored circles. Speed conditions are color-coded as follows: slow (0.7 m/s) in yellow, reference (1.05 m/s) in blue, comfortable (1.4 m/s) in orange, and fast (1.75 m/s) in purple. Top panel JND data is expressed as Weber fractions (WF; relative speed differences), revealing the expected “flat” relationship between relative JNDs and walking speed, demonstrating Weber-like scaling. The bottom panel shows the absolute JNDs from the top panel (rescaled by mean walking speed) expressed in mm/s, showing a clear positive relationship between absolute JNDs and mean walking speed.

These patterns were quantified through Just Noticeable Differences (JND_*j,c*_; Fig. 2B), which can be derived from the parameters of the logistic fits (see Methods). Consistent with Weber’s Law, relative JND values, or Weber Fraction (mean ± ste,%; expressed as a percentage of the mean walking speed) appeared constant across speeds (Fig. 2B top panel): 5.42 ± 0.665 (slow), 4.807 ± 0.291 (reference), 3.828 ± 0.31 (comfortable), 4.434 ± 0.501 (fast). In contrast, absolute JND values (Fig. 2B bottom panel) indicated that sensitivity scales as a function of the background walking speed (mean ± ste, mm/s): 37.941 ± 4.657 (slow), 50.478 ± 3.057 (reference), 53.594 ± 4.346 (comfortable), and 77.588 ± 8.760 (fast). In line with Weber’s prediction, this scaling indicates reduced sensitivity (i.e., higher absolute JND) at faster walking speeds.

### Weber’s Law scaling of sensitivity is observed in all speeds except for the comfortable speed condition

To formally test Weber’s Law, we compared the previously described logistic model in which *µ*_*j,c*_ was condition-specific 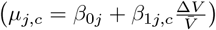 with a condition-agnostic model that assumed equal sensitivity to relative speed differences across all walking conditions, as prescribed by Weber’s Law 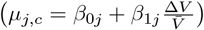. Models were compared using the difference in Akaike Information Criterion (Δ*AIC*) [25], where Δ*AIC* values higher than 6 indicate a statistical difference between models’ fits [26]. We found that the condition-specific model provided a statistically better fit of the data than the condition-agnostic model (Δ*AIC* = 6.5). Thus, sensitivity of speed differences in walking exhibits scaling, but Weber’s Law does not appear to fully capture the changes in sensitivity across all walking speeds.

The condition-specific model revealed that sensitivity adheres to Weber’s Law in participants walking at the slow and fast background walking speeds, but not at the comfortable walking speeds. We used relative sensitivity contrasts between the reference and testing conditions to quantify condition-specific departures from Weber’s Law. The sensitivity to relative speed differences was similar between the reference speed and the slow (*p* = 0.072) and fast (*p* = 0.054) test conditions, but not between the reference and the comfortable speed (*p <* 0.001; Fig. 3A). In other words, when using the reference condition as the comparison point, the sensitivity at the slow and fast walking speeds followed Weber’s scaling, whereas the comfortable walking speed did not. The distinct scaling of sensitivities across testing speeds was also observed on the absolute JND values (Fig. 3B). Paired t-test analysis indicated that absolute JNDs were significantly different between the reference speed and both the slow (*p* = 0.010) and fast walking speeds (*p <* 0.001), but not between the reference and comfortable walking speeds (*p* = 0.385). Taken together, these results suggest that Weber’s Law does not uniformly capture the scaling of perception across all speeds (Fig. 3). Specifically, while the measured relative sensitivity was similar for the slow, reference and fast speeds, at the comfortable walking speed we observed increased relative sensitivity.

**Fig 3.**
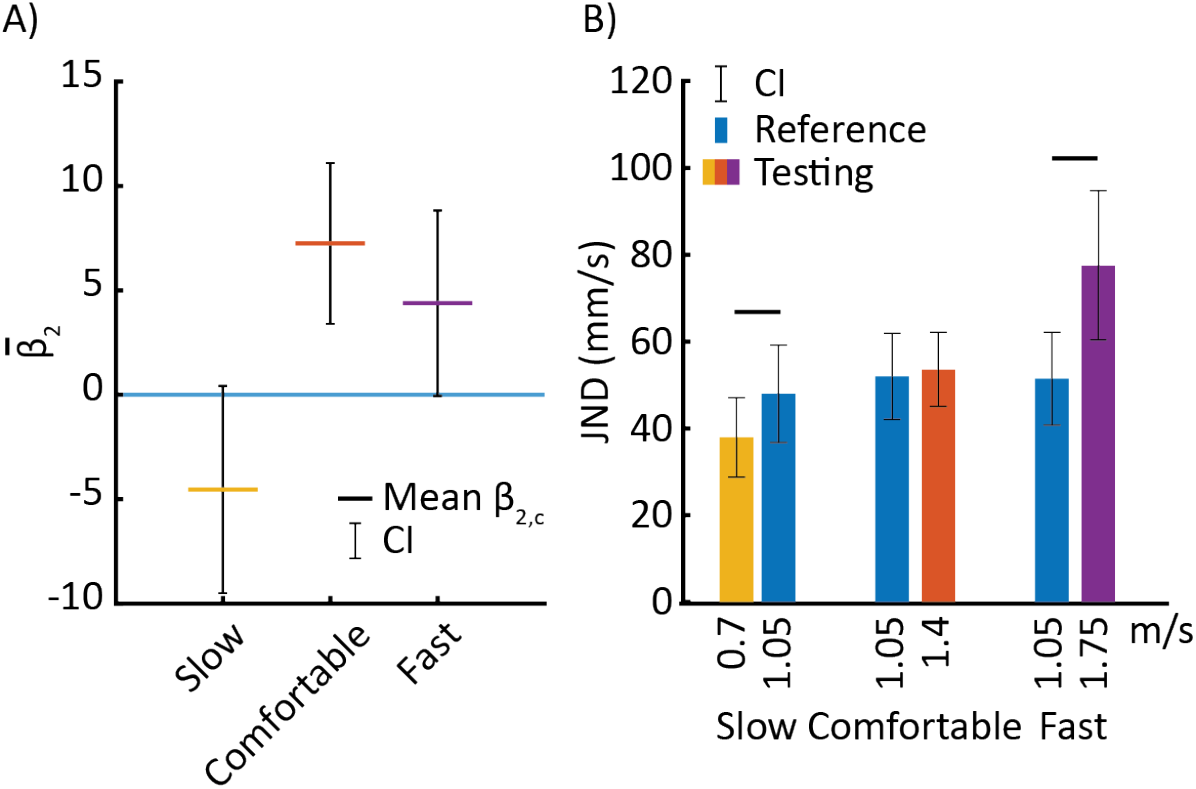
Comparison of sensitivity scaling across walking speeds. Colors denote the testing condition: slow (0.7 m/s) in yellow, comfortable (1.4 m/s) in orange, and fast (1.75 m/s) in purple. **A)** Distribution of the interaction parameter 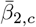, reflecting changes in relative sensitivity between the testing and reference conditions. Error bars represent 1.96 *× SE* (approximate 95% confidence intervals); bars not crossing zero indicate a significant difference in relative sensitivity. Under Weber’s Law, 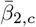 values would not differ significantly from zero (error bars crossing zero). **B)** Each bar represents the group-level average JND (in mm/s; absolute speed difference sensed 75% of the time) for a given speed condition (color-coded as in other panels). Error bars indicate 95% confidence intervals. Horizontal black lines between bars denote statistically significant differences based on paired t-tests across testing and reference conditions. The significance level for the tests was set to *α* = 0.0167.

### JND estimates derived from drift-diffusion model capture Weber-like scaling of sensitivity to speed differences

Drift-Diffusion models (DDM) are a mechanistic framework for decision-making that describes how participants accumulate noisy evidence over time until a choice is made. We have previously used this framework to predict people’s perceptual responses solely based on reaction times (i.e., time to respond after the start cue) [20]. Here, we tested the robustness of this approach across different walking speeds to assess whether this framework for evidence accumulation could mechanistically explain Weber’s scaling of sensitivity to speed differences in walking. First, we found that DDM could fit the non-linear relation between reaction times and speed differences in all walking speeds (Fig. 4A). Importantly, in our DDM implementation, the bias and sensitivity parameters (*β*_0*j,c*_ and *β*_1*j,c*_) correspond directly to those used in our primary analysis. This allowed us to predict JND_*j,c*_ values for individual subjects (*j*) and conditions (*c*) from DDM parameters fitted solely to reaction time data. The JND_*j,c*_ values derived from the DDM indicated that sensitivity to absolute speed differences scaled as a function of mean walking speed (Fig. 4B), with participants walking slower exhibiting greater sensitivity (smaller JNDs) than those walking faster (larger JNDs). Paired t-tests showed significant differences between slow and reference speeds ( *p <* 0.001), and between reference and fast speeds (*p* = 0.002), but not between the comfortable and reference speeds (*p* = 0.021) after Bonferroni correction (*α* = 0.0167). These results match those of our primary analysis, indicating that the DDM is able to capture the scaling of sensitivity to speed differences in walking from reaction time data alone.

**Fig 4.**
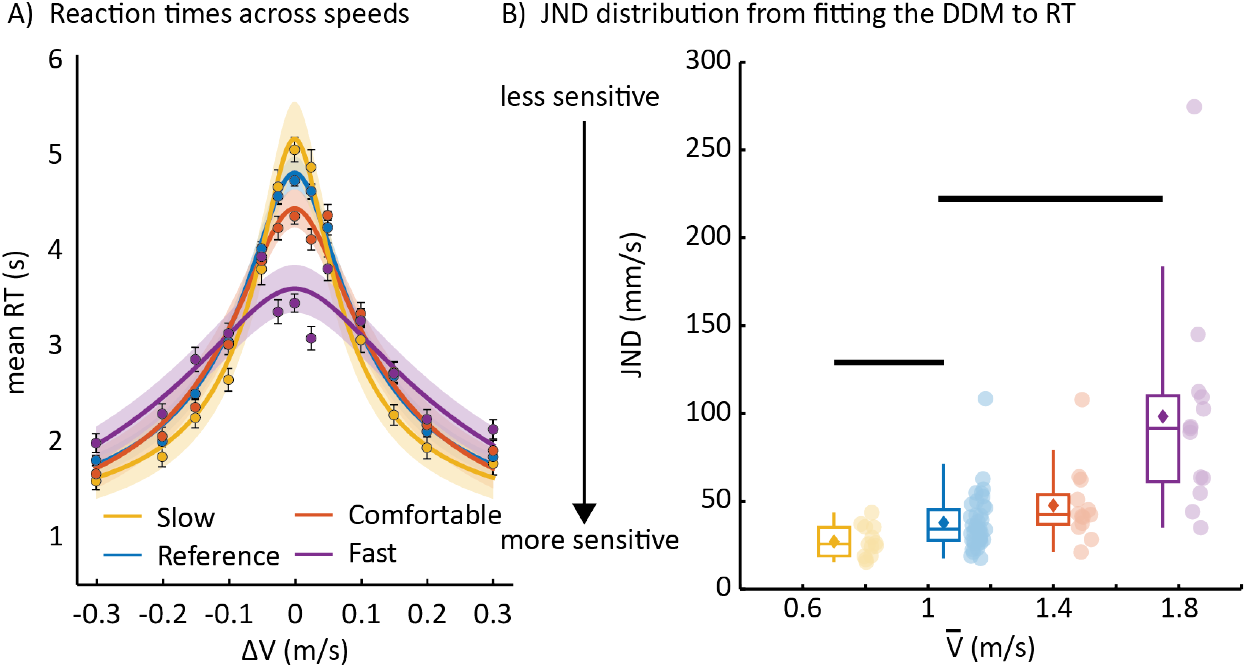
Sensitivity analysis using reaction time data alone through a drift-diffusion model (DDM). Reference speed data are pooled across participants in all three groups. **A)** DDM fits to reaction times (RTs) for all conditions/speeds. The x-axis represents absolute stimulus magnitude (Δ*V*), and the y-axis shows mean reaction time in seconds. Individual fits for each condition are summarized as thick lines (average) and shaded areas (standard error). Colors denote the speed conditions: slow (0.7 m/s), reference (1.05 m/s), comfortable (1.4 m/s), and fast (1.75 m/s). **B)** Absolute JNDs derived from the RT fits shown in panel **A**, displayed as boxplots. Means are indicated by diamonds, and individual participants by circles. Colors correspond to the speed conditions shown in panel **A**. Horizontal black lines between boxplots denote statistically significant differences based on paired t-tests between testing and reference conditions. The significance level was set to *α* = 0.0167.

## Discussion

Our primary findings demonstrate that perceptual sensitivity to speed differences during walking follows Weber’s Law. As predicted, participants were more sensitive (i.e., had lower JNDs) when walking at slower speeds compared to faster speeds. When expressed as Weber fractions, sensitivity remained approximately constant ( 4.4%–5.8%) across walking speeds (from 0.7 to 1.75 m/s), indicating that relative speed discrimination is preserved across a broad range of mean walking speeds. Importantly, we observed a deviation from Weber’s Law at comfortable walking speeds (1.4 m/s), where relative sensitivity (Weber fraction of 3.8%) was higher than what Weber’s Law predicts. This enhanced sensitivity at habitual walking speeds suggests a functional specialization of the perceptual system for ecologically relevant contexts. Finally, our drift-diffusion model analysis of reaction times validates these findings and extends this mechanistic framework explaining Weber’s Law in static, uni-sensory tasks to dynamic, multi-sensory motor tasks, such as walking.

### Weber’s Law observed in speed perception during walking

Perceptual sensitivity during walking is consistent with Weber’s Law, a principle observed across sensory modalities assessed with uni-sensory and multi-sensory perceptual tasks [8, 27], and suggests that similar neural mechanisms govern somatosensory processing during multi-sensory motor behaviors such as locomotion. The nervous system appears to maintain relative sensitivity to speed perturbations across the range of functional walking speeds, supporting effective detection and correction of gait asymmetries. By preserving an approximately constant Weber fraction, the perceptual system ensures that the threshold for detecting asymmetries scales appropriately with walking speed, allowing sufficient sensitivity to correct meaningful deviations without overreacting to minor, inconsequential differences. At faster walking speeds, for example, small asymmetries may be less relevant or too costly to correct continuously, whereas at slower speeds, heightened sensitivity may facilitate precise control. Even subtle inter-limb speed differences can disrupt coordination and reduce energetic efficiency [28, 29], highlighting the importance of flexible perceptual tuning. This adaptive scaling likely balances stability and efficiency, adjusting perceptual thresholds to match the behavioral demands of the walking context.

### Speed-specific departures from Weber’s Law reveal functional specialization

While our data generally support Weber’s Law, we found that sensitivity at the comfortable walking speed (1.4 m/s) deviated from Weber’s scaling. Notably, in a subset of participants from this study, we measured comfortable walking speed and the mean speed (1.35 ± 0.16 m/s) was even closer to the comfortable condition than to the reference condition. Specifically, instead of a constant Weber fraction across walking speeds, we observed a smaller Weber fraction at the comfortable walking speed than at other walking speeds. This indicates that perceptual sensitivity at this frequently encountered velocity was higher than expected under Weber’s Law. Such enhanced sensitivity at habitual walking speeds may provide selective advantages for detecting gait perturbations during everyday locomotion. In other words, the nervous system may have evolved specialized mechanisms for processing speed-related information at ecologically relevant walking velocities, prioritizing the detection of asymmetries most likely to be encountered during natural walking.

Several mechanisms may contribute to the enhanced sensitivity at habitual walking speeds. First, familiarity-based processing may play a crucial role. The influence of stimulus familiarity on sensory processing is well-established in visual neuroscience, where familiar stimuli produce sharpened neuronal response dynamics [30] and maintain heightened responsiveness to subsequent sensory inputs [31]. This responsiveness to new and salient sensory input captures attention and facilitates its encoding [32], creating sustained readiness for detecting perturbations within familiar contexts. By analogy, walking at a habitual speed may engage perceptual and motor systems that are finely tuned through repeated experience, allowing the nervous system to detect smaller speed asymmetries than expected by Weber’s Law. This enhanced discrimination likely reflects optimized neural tuning and processing efficiency within familiar contexts, an adaptation that prioritizes precise error detection during the speeds most commonly used in everyday locomotion.

Our observation of enhanced sensitivity at comfortable walking speeds aligns with evidence of similar departures from Weber’s law across sensory modalities at ecologically relevant stimulus ranges. Computational studies have shown that, when expressed as Weber fractions, sensitivity metrics can follow convex patterns across stimulus intensities during pure tone and light brightness discrimination, with peak discrimination efficiency occurring at intermediate ranges before declining at both low and high intensities [33, 34]. In vestibular self-motion perception, discrimination thresholds saturate rather than scale proportionally at higher amplitudes commonly experienced during natural activities, showing better performance for naturally occurring stimulus ranges [35, 36]. These patterns suggest that sensory systems exhibit functional specialization within the stimulus ranges most frequently encountered during everyday behavior, maintaining enhanced sensitivity where it is most behaviorally relevant. In auditory frequency discrimination, listeners exhibit peak sensitivity at approximately 2000 Hz [37, 38], where human hearing is optimally tuned for speech intelligibility [39], with discrimination performance varying systematically across the frequency spectrum rather than following strict Weber scaling. Analogous to our finding of a sensitivity plateau at comfortable walking speeds, frequency discrimination shows minimal just-noticeable differences within this ecologically critical range, with performance declining at both lower and higher frequencies. These patterns suggest that sensory systems exhibit functional specialization within the stimulus ranges most frequently encountered during everyday behavior, maintaining enhanced sensitivity where it is most behaviorally relevant.

### Drift-diffusion modeling validates findings and extends computational frameworks to dynamic tasks

Our drift-diffusion model analysis of reaction times complements and enhances our choice-based results, indicating that a single evidence accumulation process can effectively account for both choice patterns and reaction time distributions in a dynamic perceptual task, such as walking. Most classical DDM studies focus on brief perceptual judgments, typically with reaction times in the range of hundreds of milliseconds [23, 40]. In contrast, our walking task involves decision intervals extending up to ten seconds. This demonstrates that the underlying principles of evidence accumulation can generalize to longer, more dynamic contexts, as we have shown previously [20]. Notably, our analysis of reaction times revealed Weber-like scaling of perceptual sensitivity across varying background walking speeds, marking what we believe is the first demonstration of this relationship in a locomotor perceptual task.

While only a limited number of studies have linked the structure of reaction times to Weber’s Law, Pardo-Vazquez and colleagues demonstrated that in rats, reaction time distributions scale with stimulus magnitude in a way that is consistent with Weber’s Law [8]. Here, we extend this idea to human locomotion. By applying standard DDMs to our reaction time data, we show that the same patterns emerge, suggesting that Weber’s Law arises from proportional modulation of drift rates, likely reflecting speed-specific neural gains.

Although rats and humans differ in the complexity of their decision-making mechanisms, both show consistency with Weber’s Law, implying that proportional encoding of stimulus differences may be a general property of perceptual systems. In humans, higher-order cognitive functions modulate strategy-driven decision-making processes [41–44]. However, this flexibility in the accumulation process does not disrupt the basic scaling relationship between stimulus magnitude and perceptual sensitivity.

Our reaction time approach demonstrates the feasibility of estimating perceptual sensitivity during walking without choice data. Future work could enhance this framework by incorporating lapse rate parameters for both choice and reaction time analyses [45, 46] and by exploiting full reaction time distributions rather than means alone. These refinements may enhance cross-method consistency and provide a more comprehensive characterization of evidence accumulation dynamics in locomotor tasks. Future studies should explore how aging, neurological conditions, or interventions may alter these processes during locomotion.

## Clinical implications

Our findings provide new insights for locomotor adaptation research and rehabilitation practices. The observation that perceptual sensitivity to inter-limb speed differences is enhanced at comfortable walking speeds suggests that perceptual training protocols could be optimized by targeting habitual speed ranges. In clinical contexts, gait asymmetry detection training may be most effective when performed at these comfortable speeds, where the nervous system shows specialized processing and heightened discrimination capabilities.

Moreover, this work has direct relevance for gait asymmetry detection in neurological populations. Gait asymmetries at comfortable walking speeds are common markers of conditions such as post-stroke hemiparesis and are linked to long-term musculoskeletal degeneration, inefficient walking and fall risk [47, 48]. Impaired or altered perceptual scaling may contribute to the persistence of asymmetrical gait patterns following neurological injury. Enhancing perceptual sensitivity at these familiar speeds, through targeted perceptual-motor training or augmented feedback, could therefore improve error detection and facilitate more effective locomotor adaptation in clinical populations.

Additionally, the systematic characterization of speed discrimination thresholds across a broad range of walking speeds provides a foundation for developing objective assessment tools for locomotor impairments. Deviations from these patterns (i.e., reduced relative sensitivity at certain speeds) could serve as a clinical marker of sensorimotor impairments. Such markers may offer a more nuanced assessment than gait speed alone, potentially enabling earlier detection and more targeted interventions than current clinical assessments allow.

## Conclusion

In summary, we show that perceptual sensitivity to inter-leg speed differences during walking conforms broadly to Weber’s Law, with proportional scaling of discrimination thresholds across speeds. However, enhanced sensitivity at comfortable walking speeds indicates a functional tuning of perceptual mechanisms to the locomotor conditions most relevant to human behavior. Reaction time–based modeling corroborates these findings, extending decision making theoretical frameworks to the study of continuous, dynamic perceptual processes. These results provide a foundation for developing more effective assessment and intervention strategies for locomotor impairments while contributing to our broader understanding of how sensory processing supports adaptive motor control during human locomotion.

Future studies should also explore the neural mechanisms underlying the speed-specific departures from Weber’s Law that we observe. Neuroimaging or electrophysiological approaches could investigate whether enhanced processing at comfortable walking speeds reflects specialized cortical or subcortical mechanisms. Additionally, longitudinal studies examining how speed perception changes with locomotor training or following neurological injury could provide insights into the plasticity of these perceptual mechanisms.

## Data Availability

Data and code are available at [GitHub/OSF link to be provided].

## Materials and methods

### Data Collection

#### Participants

Thirty-nine neurotypical young adults (22 females, 23.8 ± 5.4 y.o.) participated in this study. Four of the participants reported left-handedness, and two reported being left-footed when kicking a ball [49]. Participants were randomly assigned to one of three groups: Slow (n=13), Fast (n=13), or Comfortable (n=13), corresponding to mean walking speeds slower, faster, or within the typical range for healthy young adults [24]. The reference speed condition, which was common to all groups, was used in prior work [20]. All participants were naïve to the experimental protocol and had never experienced split-belt walking. The study protocol was approved by the University of Pittsburgh’s Institutional Review Board (IRB) in accordance with the Declaration of Helsinki, and all participants gave written informed consent prior to participation.

#### Perceptual task

We measured sensitivity to inter-leg speed differences using a two-alternative forced choice (2AFC) task (See Fig. 1B). The task began with participants walking with both legs moving at the same speed, followed by an abrupt transition to a belt speed difference (stimulus: Δ*V* = *V*_*R*_ − *V*_*L*_). The stimulus was introduced by speeding up one belt and slowing down the other, sequentially and in equal amounts, so that the mean belt speed was constant throughout. Each belt’s speed was updated when the corresponding foot was in the swing phase, starting with the right belt. An auditory cue signaled the start of the response window (gray shaded area in Fig. 1B and C), which lasted 8 strides for participants in the Slow and Comfortable speed groups, and 9 seconds for the Fast group. During this period, participants reported which leg was perceived to be moving more slowly by pressing a button on a handheld controller. The perceptual task ended once a response was recorded or if they failed to respond by the end of the response window.

All auditory cues were delivered through noise-canceling headphones, including trial start, end, and warning cues. After each response, participants heard a confirmation message on their selection (e.g.,”Left is slow”). To ensure that participants completed the trial based solely on somatosensory perception, they listened to white noise and wore a drape that occluded their view of their feet, masking any external auditory or visual cues that could influence perceptual responses.

#### Testing protocol

The experimental protocol is summarized in Fig. 1A. Each participant completed two sessions of data collection on separate days, typically one day apart. Each session included two average walking speed conditions: a “reference” condition (1.05 m/s, common to all participants) and a group-specific “testing” condition (0.7 m/s for Slow, 1.4 m/s for Comfortable, or 1.75 m/s for Fast). These speeds were chosen to span a broad range: the slow and fast speed conditions deviated substantially from typical walking speeds for healthy young adults, while the reference and comfortable speeds were closer to natural walking behavior ( [24, 50, 51]). For a subset of participants from this study (n=12), we also measured comfortable walking speed using a standard 10-meter walk test (10MWT). This subset showed a mean preferred walking speed of 1.35 ± 0.16 m/s, which was even closer to the comfortable experimental condition (1.4 m/s) than to the reference condition (1.05 m/s). This value is consistent with our previous observations in age-equivalent healthy populations performing the same assessment.

Each session began with a familiarization period, where participants experienced 4 practice trials per speed condition. During this phase, participants received both visual feedback (live graphic displaying the speeds of each belt) and verbal guidance from the experimenters to ensure participants were comfortable with the procedure before data collection. Formal data collection followed, consisting of 3 blocks of 56 trials each. The speed conditions were interleaved by block, and the first speed experienced was randomized across participants to counterbalance any possible effects of experiencing a slow or fast speed first. Within a block, each non-zero stimulus magnitude ( ±0.025, ±0.05, ±0.1, ±0.15, ±0.2, and ±0.3 m/s) was presented 4 times, while null trials (0 m/s) were presented 8 times in pseudo-random order. The order was maintained across participants. The treadmill was stopped between blocks, and once within them, allowing for 3-minute breaks to avoid mental and physical fatigue. The first perceptual trial after starting the treadmill was omitted from the analysis, as in [20], but the same stimulus magnitude was collected at the end of the block to ensure the same number of repetitions per stimulus. This was done to avoid a lack of preparedness for the perceptual task. Across all participants, we collected a total of *M* = 234 blocks and analyzed a total of 13, 104 perceptual trials (56 perceptual tasks *×* 6 experimental blocks *×* 39 participants). Non-response trials were excluded from the analysis (a total of 37 out of 13,104 trials), corresponding to 0.28% of all trials. These occurred at various speed differences: 13 trials at 0 mm/s, 10 at 25 mm/s, 7 at 50 mm/s, 5 at 150 mm/s, and 2 at 300 mm/s.

### Data Analysis

#### Psychometric curve fitting

We hypothesized (*H*_0_) that relative perceptual sensitivity is invariant across mean walking speeds, consistent with Weber’s Law. In this case, the slope of the psychometric function as a function of relative stimulus size 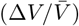 should remain constant across conditions. The alternative hypothesis (*H*_*A*_) was that sensitivity differs across walking speeds, indicating a deviation from Weber’s Law.

To test this hypothesis, we modeled participants’ binary choices (left or right) in the 2AFC tasks using a logistic psychometric function:

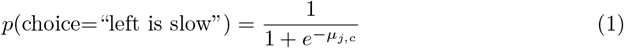

We selected a two-component linear model for *µ*_*j,c*_, with a bias and a sensitivity term, as in [20]:

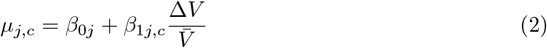

Here, *j* refers to individual participants and *c* to walking speed conditions (i.e., slow, reference, comfortable, or fast). The intercept *β*_0*j*_ captures participant specific response bias (a tendency to favor one leg over the other), while the slope *β*_1*j,c*_ quantifies sensitivity, how strongly sensory stimuli influence perceptual judgments under each condition.

For convenience in the analysis, and exploiting the existence of a common reference condition experienced by all participants, the sensitivity was expressed as:

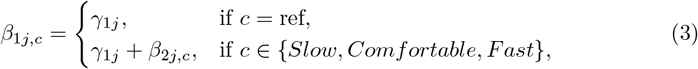

Specifically, under the reference condition, the sensitivity term is given by *γ*_1*j*_, where *γ*_1*j*_ represents participant *j*’s baseline sensitivity to relative speed differences 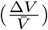. For any testing condition (*c* ≠ ref), sensitivity is expressed as *γ*_1*j*_ + *β*_2*j,c*_. The parameter *β*_2*j,c*_ therefore quantifies the difference in sensitivity between the reference and the testing condition for participant *j*. A positive *β*_2*j,c*_ indicates increased sensitivity during the testing condition relative to the reference, while a negative *β*_2*j,c*_ indicates reduced sensitivity. Results from the model are displayed in Table 1.

**Table 1.**
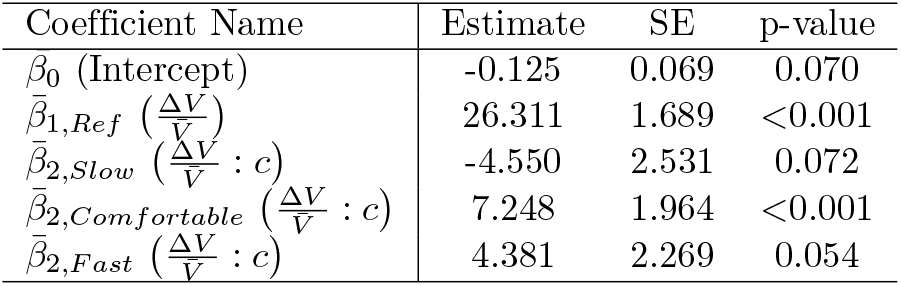
Model estimates from Eq. 2 and p-values for each mean walking speed.

Parameters were estimated using a hierarchical generalized linear mixed-effects model (GLMM) implemented in MATLAB’s *fitglme* function with maximum likelihood estimation via Laplace approximation. The model treated participant as a random effect to account for individual differences in both bias and sensitivity, relative stimulus magnitude 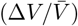 as a continuous predictor, and walking speed condition (slow, reference, comfortable, or fast) as a categorical variable. This hierarchical approach allows individual estimates to be informed by the full dataset, with population-level parameters serving as regularization for individual-level estimates. This method provides less variable inferences for individual parameters compared to fitting separate curves for each participant and condition [52].

### Just noticeable difference estimation

We also reported sensitivity using the just noticeable difference (JND), defined as the smallest reliably detectable belt speed difference, which is a more intuitive and widely used metric in psychophysics. This alternative representation facilitates comparisons with classic findings and helps interpret sensitivity changes in the context of Weber’s Law. JND is mathematically derived from the slope of the fitted psychometric curve for each subject as shown in Eq. 4 [53].

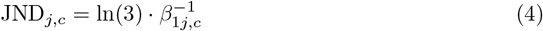

To facilitate interpretation, we report psychometric functions and JNDs in both relative and absolute magnitude terms. Relative values express speed differences as a proportion of mean walking speed 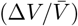, whereas absolute values (mm/s) were obtained by multiplying the relative difference by the mean walking speed for each condition.

### Drift Diffusion Modeling

Our two-alternative forced choice task recorded both participants’ binary choice responses and reaction times (RT), which can offer additional insight into the underlying decision process. Previous work has demonstrated that reaction times alone can provide estimates of perceptual sensitivity (JND) in psychophysical tasks [20]. To test the robustness of our findings, we implemented a drift diffusion model (DDM) analysis to determine whether the scaling patterns observed in our choice data could be reproduced using reaction time data alone. This approach allows us to evaluate whether the DDM framework can capture Weber-like scaling in walking tasks independent of the binary choice responses.

In our task, we defined RT as the time elapsed between the onset of the start cue (presented after the full speed difference was set) and the participants’ response. We model the reaction time data using a simplified DDM.

The DDM is a decision-making model where participants accumulate noisy evidence (*x*(*t*)) over time *t* until a decision threshold is crossed. Evidence accumulation begins after a non-decision period (*t*_*nd*_) and evolves according to a stochastic differential equation, as observed in Eq. 5.

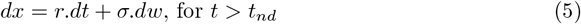

Where *r* is the drift rate, *σ* is the noise standard deviation, and *dw* ∼ *N* (0, *dt*) is Gaussian noise. Decisions occurs when the evidence reached one of two symmetric and fixed decision boundaries starting from an unbiased initial point (*x*(*t*) = 0, *t* ≤ *t*_*nd*_). The duration of evidence accumulation is referred to as the decision time (*t*_*d*_), and the sum of the decision time and non-decision times gives the overall reaction time, which is the only directly observable time parameter. This simple DDM is characterized by four parameters: the decision barrier location *a*, which quantifies the necessary evidence to reach a decision; the diffusion rate (*σ*); the drift rate (r); and the non-decision time (*t*_*nd*_), accounting for processes like stimulus encoding and motor response. This parametrization is redundant, as a constant scaling of *a, σ*, and *r* results in the same model predictions. Thus, without loss of generality, we set *a* = 1 as in [20].

In this model, the probability of the process reaching a specific decision barrier (e.g., the probability that a left choice is made, denoted *p*_*L*_) and the mean decision time have closed-form expressions, whose derivation can be found in [22, 54]. We adapted the expressions to our parametrization of the problem and obtained the results shown below. The probability of reaching the other decision barrier is defined as *p*_*R*_ = 1 − *p*_*L*_.

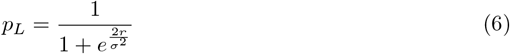

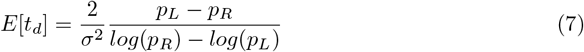

We assume that the non-decision time (*t*_*nd*_) and noise (*σ*) are stimulus-independent [55], while the drift rate *r* varies linearly with the sensory evidence (absolute speed difference, Δ*V*). The simplest such relationship is a linear one, which we choose to parametrize as:

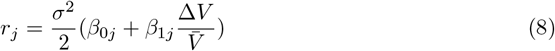

It can be observed that the psychometric curve expected from this DDM is equivalent to the ones used earlier to model perceptual choices, with 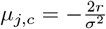 Consequently, the parameters *β*_0*j*_ and *β*_1*j*_ have the exact same interpretation as in the psychometric curve fitting approach, as a bias and sensitivity term respectively.

Model fitting was performed separately for each experimental group and speed condition using a least-squares approach on mean reaction times across stimulus magnitudes. No information about choices was used. The model to be fitted then has four scalar degrees of freedom: *t*_*nd*_, *σ, β*_0_, and *β*_1_. Since the DDM yields both the bias term (*β*_0*j*_) and the sensitivity term (*β*_1*j*_), we can derive the JND directly from these estimates. This allowed us to evaluate whether JNDs scale proportionally with walking speed, consistent with Weber’s Law, based solely on reaction time behavior. By doing so, we directly test whether the DDM captures Weber-like scaling in walking tasks (as shown in our main primary behavioral analysis) and examine the extent to which this modeling framework accounts for perceptual sensitivity in locomotor contexts. Full model derivation and implementation details are available in [20].

### Sample size estimation

. We determined the required sample size using a simulation-based power analysis performed in MATLAB, based on unpublished psychometric parameters from a previous study [56]. Our goal was to estimate the minimum number of participants required to detect differences in perceptual sensitivity (i.e., the slope of a logistic function) between two speed conditions, referred to as “reference” and “testing.” Under the null hypothesis, we assumed that the slopes of the psychometric curves in both conditions were equal, indicating similar perceptual sensitivity (i.e. no scaling of sensitivity with mean walking speed). Under the alternative hypothesis, we assumed that perceptual sensitivity varied between conditions due to differences in speed, leading to distinct slopes that are proportional to mean walking speed, consistent with Weber’s Law.

We used logistic regression to model binary choices as a function of belt-speed difference (Δ*V*). In the simulation, we drew participant-specific slopes from a set of empirical estimates obtained from [56]. For each participant, we simulated binary responses at our tested Δ*V*, with 12 repetitions per level, for both speed conditions. For each condition, we scaled the stimulus magnitude of the logistic function with the condition mean walking speed (reference: 1.05, testing: 0.7, 1.4, or 1.75 m/s). A total of 1000 simulation runs were conducted per sample size. In each run, we generated synthetic responses for a given number of subjects and tested whether the confidence intervals (CIs) of the group slope estimates for each condition overlapped. The proportion of runs in which the CIs for the two conditions did not overlap was used as an empirical estimate of power. The simulations showed that a sample size of 13 participants was sufficient to achieve over 80% power to detect a significant difference in slopes between speed conditions. We used the largest required sample size across all conditions to ensure consistent power.

Later in the study, our data analysis strategy evolved to use logistic linear mixed-effects models (GLMMs) instead of fitting each condition independently. This fitting method has been shown to provide less variable inferences for individual estimates given that is assumes that individual parameters are drawn from underlying Gaussian distributions [52], serving as a regularization in the fitting procedure and accounting for the nested structure in out dataset.

## Statistical analysis

To test for significant differences in sensitivity across speed conditions, we used the 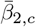 estimates from the GLMM fits (see Psychometric Curve Fitting), which quantify the deviation in sensitivity from the reference condition (i.e., the interaction effect). Statistical significance was assessed using the Wald test for the fixed effect 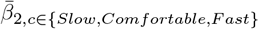, which evaluates if the model coefficient differs significantly from zero (*H*_0_ : *β* = 0). The significance level for the tests was set to *α* = 0.05.

Lastly, we performed paired t-tests comparing absolute JNDs between the testing and reference conditions within each of the three experimental groups (Slow, Comfortable, and Fast). These JNDs were derived from two independent sources: (1) the generalized linear mixed model (GLMM) used in our primary analysis of choice behavior, and (2) the drift diffusion model (DDM) fit to reaction time data. To correct for multiple comparisons, we applied a Bonferroni correction, resulting in an adjusted significance threshold of *α* = 0.0167.

## References

1. Shadmehr R, Mussa-Ivaldi FA. Adaptive representation of dynamics during learning of a motor task. Journal of Neuroscience. 1994;14(5):3208–3224. doi:10.1523/JNEUROSCI.14-05-03208.1994.

2. Smith MA, Ghazizadeh A, Shadmehr R. Interacting adaptive processes with different timescales underlie short-term motor learning. PLOS Biology. 2006;4(6):e179. doi:10.1371/journal.pbio.0040179.

3. Taylor JA, Ivry RB. Flexible cognitive strategies during motor learning. PLOS Computational Biology. 2011;7(3):e1001096. doi:10.1371/journal.pcbi.1001096.

4. Torres-Oviedo G, Vasudevan E, Malone L, Bastian AJ. Locomotor adaptation. Progress in Brain Research. 2011;191:65–74. doi:10.1016/B978-0-444-53752-2.00013-8.

5. Ganel T, Chajut E, Algom D. Visual coding for action violates fundamental psychophysical principles. Curr Biol. 2008 Jul;18(14):R599–R601. doi:10.1016/j.cub.2008.04.052.

6. Zanker JM. Does motion perception follow Weber’s law? Perception. 1995;24(4):363–372. doi:10.1068/p240363.

7. McGill WJ, Goldberg JP. A study of the near-miss involving Weber’s law and pure-tone intensity discrimination. Perception & Psychophysics. 1968;4(2):105–109. doi:10.3758/BF03209518.

8. Pardo-Vazquez JL, Castiñeiras-de Saa JR, Valente M, Damião I, Costa T, Vicente MI, Mendonça AG, Mainen ZF, Renart A. The mechanistic foundation of Weber’s law. Nat Neurosci. 2019;22(9):1493–1502. doi:10.1038/s41593-019-0439-7.

9. Deco G, Scarano L, Soto-Faraco S. Weber’s law in decision making: Integrating behavioral data in humans with a neurophysiological model. J Neurosci. 2007;27(42):11192–11200. doi:10.1523/JNEUROSCI.1072-07.2007.

10. Contu S, Marini F, Cappello L, Masia L. Robot-assisted assessment of wrist proprioception: Does wrist proprioceptive acuity follow Weber’s law? Proc Annu Int Conf IEEE Eng Med Biol Soc. 2016;2016:4610–4613. doi:10.1109/EMBC.2016.7591754.

11. Stone H, Bosley JJ. Olfactory discrimination and Weber’s Law. Percept Mot Skills. 1965;20(2):657–665. doi:10.2466/pms.1965.20.2.

12. Lauziere S, Miéville C, Duclos C, Aissaoui R, Nadeau S. Perception threshold of locomotor symmetry while walking on a split-belt treadmill in healthy elderly individuals. Percept Mot Skills. 2014;118(2):475–490. doi:10.2466/25.15.PMS.118k17w6.

13. Hoogkamer W, Bruijn SM, Potocanac Z, Van Calenbergh F, Swinnen SP, Duysens J. Gait asymmetry during early split-belt walking is related to perception of belt speed difference. J Neurophysiol. 2015;114:1705–1712. doi:10.1152/jn.00937.2014.

14. Wutzke CJ, Faldowski RA, Lewek MD. Individuals poststroke do not perceive their spatiotemporal gait asymmetries as abnormal. Phys Ther. 2015;95(9):1244–1253. doi:10.2522/ptj.20140482.

15. Vazquez A, Statton MA, Busgang SA, Bastian AJ. Split-belt walking adaptation recalibrates sensorimotor estimates of leg speed, but not position or force. J Neurophysiol. 2015;jn.00302.2015. doi:10.1152/jn.00302.2015.

16. Statton MA, Vazquez A, Morton SM, Vasudevan EVL, Bastian AJ. Making sense of cerebellar contributions to perceptual and motor adaptation. Cerebellum. 2018;17(2):111–121. doi:10.1007/s12311-017-0879-0.

17. Leech KA, Day KA, Roemmich RT, Bastian AJ. Movement and perception recalibrate differently across multiple days of locomotor learning. J Neurophysiol. 2018;120:2130–2137. doi:10.1152/jn.00355.2018.-Learning.

18. Rossi C, Roemmich RT, Bastian AJ. Understanding mechanisms of generalization following locomotor adaptation. npj Sci Learn. 2024;9(1):48. doi:10.1038/s41539-024-00258-2.

19. Rossi C, Leech KA, Roemmich RT, Bastian AJ. Automatic learning mechanisms for flexible human locomotion. eLife. 2024 Oct;13:RP101671. doi:10.7554/eLife.101671.1.

20. Gonzalez-Rubio M, Torres-Oviedo G, Iturralde PA. Characterizing human perception of speed differences in walking: Insights from a Drift Diffusion Model. eNeuro. 2025;12(5). doi:10.1523/ENEURO.0343-23.2025.

21. Ratcliff R. A theory of memory retrieval. Psychological Review. 1978;85(2):59–108. doi:10.1037/0033-295X.85.2.59.

22. Bogacz R, Brown E, Moehlis J, Holmes P, Cohen JD. The physics of optimal decision making: A formal analysis of models of performance in two-alternative forced-choice tasks. Psychological Review. 2006;113(4):700–765. doi:10.1037/0033-295X.113.4.700

23. Gold JI, Shadlen MN. The neural basis of decision making. Annual Review of Neuroscience. 2007;30:535–574. doi:10.1146/annurev.neuro.29.051605.113038.

24. Ralston HJ. Energy-speed relation and optimal speed during level walking. Int Z Angew Physiol. 1958;17(4):277–283. doi:10.1007/BF00698754.

25. Cavanaugh JE, Neath AA. The Akaike information criterion: Background, derivation, properties, application, interpretation, and refinements. WIREs Computational Statistics. 2019;11(3):e1460. doi:10.1002/wics.1460.

26. Symonds MRE, Moussalli A. A brief guide to model selection, multimodel inference and model averaging in behavioural ecology using Akaike’s information criterion. Behavioral Ecology and Sociobiology. 2011;65(1):13–21. doi:10.1007/s00265-010-1037-6.

27. Fechner GT. Elements of psychophysics. Leipzig: Breitkopf und Härtel; 1860.

28. Finley JM, Bastian AJ, Gottschall JS. Learning to be economical: the energy cost of walking tracks motor adaptation. The Journal of Physiology. 2013;591(4):1081–1095. doi:10.1113/jphysiol.2012.245506.

29. Butterfield JK, Collins SH. The energy cost of split-belt walking for a variety of belt speed combinations. Journal of Biomechanics. 2022;132:110905. doi:10.1016/j.jbiomech.2021.110905.

30. Meyer T, Walker C, Cho RY, Olson CR. Image familiarization sharpens response dynamics of neurons in inferotemporal cortex. Nat Neurosci. 2014;17(10):1388–1394. doi:10.1038/nn.3794.

31. Manahova ME, Spaak E, de Lange FP. Familiarity increases processing speed in the visual system. J Cogn Neurosci. 2020;32(4):722–733. doi:10.1162/jocn_a_01507.

32. Pedale T, Santangelo V. Perceptual salience affects the contents of working memory during free-recollection of objects from natural scenes. Front Hum Neurosci. 2015;9:60. doi:10.3389/fnhum.2015.00060.

33. Penconek M. Weber’s Law as the emergent phenomenon of choices based on global inhibition. Frontiers in Neuroscience. 2025;19:1532069. doi:10.3389/fnins.2025.1532069.

34. Freyman RL, Nelson DA. Frequency discrimination of short-versus long-duration tones by normal and hearing-impaired listeners. Journal of Speech and Hearing Research. 1987;30(1):28–36. doi:10.1044/jshr.3001.28.

35. Carriot J, Cullen KE, Chacron MJ. The neural basis for violations of Weber’s law in self-motion perception. Proceedings of the National Academy of Sciences. 2021;118(36):e2025061118. doi:10.1073/pnas.2025061118.

36. Mallery RM, Olomu OU, Uchanski RM, Militchin VA, Hullar TE. Human discrimination of rotational velocities. Experimental Brain Research. 2010;204(1):11–20. doi:10.1007/s00221-010-2288-1.

37. Benson D. Music: A Mathematical Offering. Cambridge: Cambridge University Press; 2006.

38. Robinson DW, Dadson RS. A re-determination of the equal-loudness relations for pure tones. British Journal of Applied Physics. 1956;7(166). doi:10.1088/0508-3443/7/5/302.

39. Dinh TA. Improving speech intelligibility through spectral style conversion. PhD Thesis, Oregon Health & Science University; 2021.

40. Ratcliff R, McKoon G. Drift Diffusion Decision Model: Theory and data. Neural Computation. 2008;20(4):873–922. doi:10.1162/neco.2008.12-06-420.

41. Rahnev D, Nee DE, Riddle J, Larson AS, D’Esposito M. Causal evidence for frontal cortex organization for perceptual decision making. Proceedings of the National Academy of Sciences. 2016;113(21):6059–6064. doi:10.1073/pnas.1522551113.

42. Heekeren HR, Marrett S, Bandettini PA, Ungerleider LG. A general mechanism for perceptual decision-making in the human brain. Nature. 2004;431(7010):859–862. doi:10.1038/nature02966.

43. Hanks TD, Summerfield C. Perceptual Decision Making in Rodents, Monkeys, and Humans. Neuron. 2017;93(1):15–31. doi:10.1016/j.neuron.2016.12.003.

44. Miller EK, Cohen JD. An integrative theory of prefrontal cortex function. Annual Review of Neuroscience. 2001;24:167–202. doi:10.1146/annurev.neuro.24.1.167.

45. Treutwein B, Strasburger H. Fitting the psychometric function. Perception & Psychophysics. 1999;61(1):87–106. doi:10.3758/bf03211951.

46. Wichmann FA, Hill NJ. The psychometric function: I. Fitting, sampling, and goodness of fit. Perception & Psychophysics. 2001;63(8):1293–1313. doi:10.3758/BF03194544.

47. Patterson KK, Parafianowicz I, Danells CJ, Closson V, Verrier MC, Staines WR, Black SE, McIlroy WE. Gait asymmetry in community-ambulating stroke survivors. Archives of Physical Medicine and Rehabilitation. 2008;89(2):304–310. doi:10.1016/j.apmr.2007.08.142.

48. Lewek MD, Braun CH, Wutzke CJ, Giuliani C. The role of movement errors in modifying spatiotemporal gait asymmetry post-stroke: A randomized controlled trial. Clinical Rehabilitation. 2018;32(2):87–97. doi:10.1016/j.biomaterials.2016.06.015.Tunable.

49. Kramer JF, Balsor BE. Lower extremity preference and knee extensor torques in intercollegiate soccer players. Can J Sport Sci. 1990;15(3):180–184. PMID: 2257531.

50. Knoblauch RL, Pietrucha MT, Nitzburg M. Field studies of pedestrian walking speed and start-up time. Transp Res Rec. 1996;1538(1):27–38. doi:10.1177/0361198196153800104.

51. Schimpl M, Moore C, Lederer C, Neuhaus A, Sambrook J, Danesh J, Ouwehand W, Daumer M. Association between walking speed and age in healthy, free-living individuals using mobile accelerometry–a cross-sectional study. PLoS ONE. 2011;6(8):e23299. doi:10.1371/journal.pone.0023299.

52. Gelman A, Hill J. Data analysis using regression and multilevel/hierarchical models. Cambridge: Cambridge University Press; 2006. doi:10.1017/CBO9780511790942.

53. Bush RR. Estimation and evaluation. In: Luce RD, Bush RR, Galanter E, editors. *Handbook of Mathematical Psychology*. New York: John Wiley & Sons; 1963. p. 447–456.

54. Wagenmakers EJ, Van Der Maas HLJ, Grasman RPPP. An EZ-diffusion model for response time and accuracy. Psychonomic Bulletin and Review. 2007;14(1):3–22. doi:10.3758/BF03194023.

55. Palmer J, Huk AC, Shadlen MN. The effect of stimulus strength on the speed and accuracy of a perceptual decision. J Vis. 2005 May;5(5):376–404. doi:10.1167/5.5.1.

56. Iturralde PA, Gonzalez-Rubio M, Torres-Oviedo G. High-human acuity of speed asymmetry during walking. bioRxiv. 2020 Oct 28;2020.10.28.359281. doi:10.1101/2020.10.28.359281.

